# Integrating Recombinase-Based Feedback and Feedforward Control for Optimal Resource Decoupling

**DOI:** 10.1101/2025.05.13.653855

**Authors:** Rixin Zhang, Rong Zhang, Xiao-Jun Tian

## Abstract

Resource competition disrupts circuit modularity by introducing unintended coupling between otherwise independent gene modules, thereby compromising genetic circuit function. While various control strategies have been explored, their complexity or limited efficacy has hindered broader application. Here, we present the Re-NF-FF-Controller, a recombinase-based strategy that integrates negative feedback and feedforward regulation via promoter flipping to mitigate resource competition. Computational modeling and experimental validation demonstrate that Re-NF-FF-Controller effectively reduces resource coupling, ensuring robust gene expression and modularity. Moreover, its tunability allows for performance optimization through straightforward adjustments of recombinase enzyme levels. This strategy offers a versatile and easily implementable solution for designing reliable synthetic biological systems.

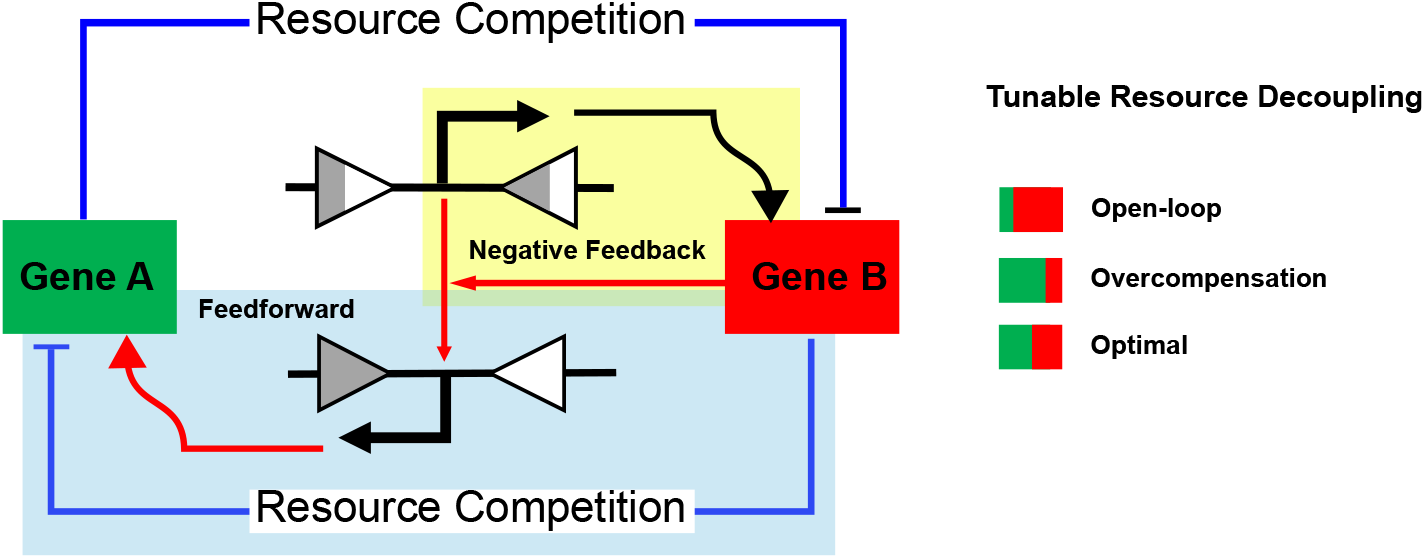

## Introduction

Synthetic biology offers a bottom-up approach for constructing genetic circuits to achieve tailored functions by leveraging previously characterized modules (1). However, emergent or unintended behaviors often arise when multiple gene modules are assembled, making the engineering process lengthy and tedious (2–5). A key challenge arises from the finite pool of cellular resources— competition for these resources can disrupt the intended function of individual modules, leading to unpredictable outcomes that deviate from the original design (6–9). This resource limitation not only introduces uncertainty but also weakens the modularity principle that underpins synthetic biology.

To restore the modularity of genetic circuits, various methods have been proposed (2, 4, 10). Using orthogonal resources can compartmentalize different gene modules into distinct groups, thereby reducing resource coupling by utilizing separate resource pools (11–13). Resource reallocation is another strategy that allows for dynamic allocation of resources to different gene modules as needed, ensuring stable expression of the desired modules and enhancing the robustness of the gene circuit (14, 15). Negative feedback and feedforward loops are also utilized for resource competition control (16–23). For example, Darlington et al. developed a resource reallocator in which a negative feedback controller was incorporated (11). By sensing changes in circuit demand, it adjusts the available ribosomes to maintain robustness. Shopera et al. introduced negative feedback into the operating system to decouple resource-coupled gene expression (24). Additionally, Barajas et al. constructed a feedforward growth rate controller by co-expressing SpoTH with a gene of interest (GOI), where SpoTH played a role in replenishing ribosomes sequestered by GOI expression (25). However, these strategies often offer limited improvements or rely on complex designs, reducing their versatility and broader applicability.

Site-specific recombinases modify DNA sequences by recognizing and acting on defined binding sites (26). Depending on the orientation of the recombinase sites, recombination can occur in three ways: integration, excision, and inversion. While recombinase-mediated integration is commonly used for DNA assembly, its excision and inversion capabilities enable precise logical control in genetic circuits. For example, recombinase has been used to construct a rewriteable, recombinase-addressable data module, which reliably records digital information by fine-tuning the synthesis and degradation rates of recombinase proteins (27). Recombinase-based negative feedback controllers have been developed to reduce gene expression variability (28). Additionally, recombinase enzymes have been employed to create a feedback controller capable of near-perfect adaptation, with three distinct operational modes that respond to external signals, taking advantage of the switching properties of recombinase enzymes (29). In this work, we present a resource controller that leverages the recombinase-mediated reversion mechanism to integrate negative feedback and feedforward control, enhancing gene circuit modularity.

## Materials and Methods

### Strains, media, and chemicals

*E. coli* strains DH5α and K-12 MG1655*ΔlacIΔaraCBAD* were utilized for circuit constructions and circuit inductions, respectively. Single DH5α colonies were inoculated into 5 ml LB broth containing either 25 μg/ml chloramphenicol, 50 μg/ml kanamycin, or 100 μg/ml ampicillin, depending on the backbones of the plasmids. For plasmid extraction, cells were grown in 15 ml tubes at 37 °C on a shaker set to 250 RPM (rotations per minute). For circuit induction and measurement, cells were induced by L-ara (L-(+)-Arabinose, Sigma-Aldrich) in a 96-well plate. L-ara was dissolved in ddH2O into a stock concentration of 25%, then diluted into appropriate working solutions for induction. The induction scheme is detailed further down in the Methods.

### Plasmids construction

Circuits were constructed in either pSB1A3, pSB1C3, or pSB3K3 backbones. The induced circuits used for measurements were on pSB3K3 (medium copy number) or pSB1C3 backbone (high copy number) in this study. The BioBricks used directly to build our circuits are listed in **Supplementary Table 1**. The sequences of other parts are listed in the **Supplementary Table 2**. The parts were first digested as designed via restriction sites for EcoRI, XbaI, SpeI, and PstI (restriction enzymes from ThermoFisher), then purified by PCR cleanup (GenElute PCR Cleanup Kit from Sigma-Aldrich) or gel electrophoresis (GelElute Gel Extraction Kit from Sigma-Aldrich) depending on the length of the products. The digested parts were ligated using T4 DNA ligase (New England BioLabs) and then transformed into *E. coli* strain DH5α. Transformants were screened by colony PCR and cultured overnight. Finally, the plasmids were extracted using GenElute Plasmids Miniprep Kit (Sigma-Aldrich) and further validated by digesting EcoRI and PstI restriction sites. Details of circuit design can be found in **Supplementary Table 3**. All sequences of plasmids were verified by whole plasmid sequencing.

### Circuit inductions

One colony of *E. coli* strain K-12 MG1655*ΔlacIΔaraCBAD* was inoculated into 300 μl LB medium containing either 50 μg/ml kanamycin (LBK) or 25 μg/ml chloramphenicol (LBC), depending on the plasmid backbone, and grown in a 5 ml culture tube for 5 hours at 37 °C on a shaker set to 250 RPM. Then, 5 μl cell culture was added into each well of the 96-well plate containing 200 μl LBK or LBC with 3 replicates of each sample. 25% L-ara was diluted into a gradient of 8.3%, 5%, 2.5%, 0.625%, 0.25%, 0.125% and 0% to prepare the working solutions, then further diluted 1000-fold into LBK or LBC to induce the gene circuits. The cells were induced overnight in a shaker and then measured on a plate reader.

Synergy H1 Hybrid Reader from BioTek was used to measure the fluorescence intensity. After adding different concentrations of inducers according to the experimental design, the assembled plates were incubated overnight at 37 °C in a shaker set to 250 RPM to allow gene expression to reach a steady state. Following incubation, 200 μl cell culture was added into each well of the 96-well plate, and the same amount of LBK or LBC without cells was used as a blank control. The optical density (OD) of the culture was measured by absorbance at 600 nm; GFP was detected by excitation/emission at 485/515 nm; RFP was detected by excitation/emission at 546/607 nm. CFP was detected by excitation/emission at 438/485 nm. To evaluate cellular burden, cell growth was monitored dynamically by measuring OD. Instead of overnight culturing in a shaker, the assembled plate was incubated in a plate reader with orbital shaking at 807 CPM (circles per minute) at 37 °C. OD was recorded every 20 minutes.

### Time-Course Analysis of Promoter Flipping

Plasmids carrying circuits C53 and C55 were transformed into *E. coli* K-12 MG1655ΔlacIΔaraCBAD. A single colony was inoculated into 2 mL of LBK medium and cultured overnight at 37 °C with shaking at 250 RPM. Circuits C53 and C55 were induced with 5 × 10^−3^% L-arabinose at 0, 3, 6, 9, and 12 hours. Finally, plasmids were extracted using the GenElute Plasmids Miniprep Kit (Sigma-Aldrich). Plasmids that underwent flipping into the attBP configuration contained the sequence from the Pbad promoter to the GFP gene, which was amplified by PCR using the primers: Forward: 5’-ccgcttctagagacattgattatttgcacggcgtcacactttg-3’ and Reverse: 5′-ggaacaggtagttttccagtagtgc-3’. Plasmids that remained in the attLR configuration contained the sequence from the Pbad promoter to the RFP gene, amplified using the same forward primer and a different reverse primer: 5’-tctggaattcgcggccgcttctagagttattaagcaccggtggagtg-3’. PCR with 16 cycles was performed to compare product concentrations at different time points.

### Microscopy

To test the flipping efficiency of integrase, gene circuits C5 and C6 were induced with 6.25 × 10^−4^% and 5 × 10^−3^% L-ara, following a protocol similar to the one described above, except that after the 5-hour incubation step, 2 μl of cell culture was added to a 15 ml culture tube containing 2 ml LBK. Then, the cells were induced overnight for observation. 2 μl of cell culture was placed at the center of a 2.4 × 5 cm No. 1 coverslip and covered with a piece of agarose gel made with PBS. Phase, GFP and RFP images were taken with a 100X oil objective on a Nikon Eclipse Ti inverted microscope (Nikon) equipped with an LED-based Lumencor SOLA SE.

### Flow cytometry

All samples were analyzed using Stratedigm S1000EON Flow Cytometer with excitation/emission filters 480 nm/530 nm (FL1-A) for GFP detection and 480 nm/>670 nm (FL3-A) for RFP after overnight induction. Each sample was represented by three biological replicates, with 10,000 events recorded per replicate for analysis. The generated data was further analyzed using MATLAB (R2024b, MathWorks).

### Coupling Index (CI)

To quantify the coupling between GFP and RFP expression, we defined a coupling index (CI) as the average deviation of the experimental GFP-RFP response curve from an ideal flat line representing complete resource decoupling. GFP levels were normalized to their baseline expression without RFP module induction, and RFP levels were normalized to their maximum expression. Experimental data were fitted using a Piecewise Cubic Hermite Interpolating Polynomial (PCHIP), and CI was computed as the average difference between the fitted curve and the ideal value of 1. For simulations, the density of data points allowed direct CI estimation without fitting. A CI of 0 indicates perfect decoupling, while negative or positive values reflect negative or positive coupling, respectively

### Mathematical models

Mathematical models based on ordinary differential equations (ODEs) were developed to describe and analyze system dynamics without a controller, as well as with the Re-NF-Controller or Re-NF-FF-Controller. Steady-state simulations were performed to assess GFP expression dependence on the RFP module across all scenarios. Further details are provided in the Supplementary Information.

## Results

### Design of Recombinase-Based Control Strategy for Minimizing Resource Coupling

In this study, we introduce a recombinase-based control strategy to minimize resource coupling between two seemingly unconnected modules in the cellular environment. For simplicity, two genes are designed in the synthetic gene circuit, green fluorescent protein (GFP) driven by a constitutive promoter and red fluorescent protein (RFP) controlled by an inducible promoter (Fig.1A). In an ideal scenario with unlimited cellular resources, these two genes should be expressed independently. Specifically, GFP expression level should remain unchanged regardless of RFP expression level, resulting in a flat interdependence curve (red line, **Fig.1B**). However, cellular resources are inherently limited, resulting in significant competition between the two genes and unintended coupling. Upon induction, the RFP-expressing module sequesters a fraction of resources, leading to a reduction in GFP expression. This resource-driven interference is reflected as a negative correlation between the expression of the two genes (blue line, **Fig. 1B**).

**Figure 1.**
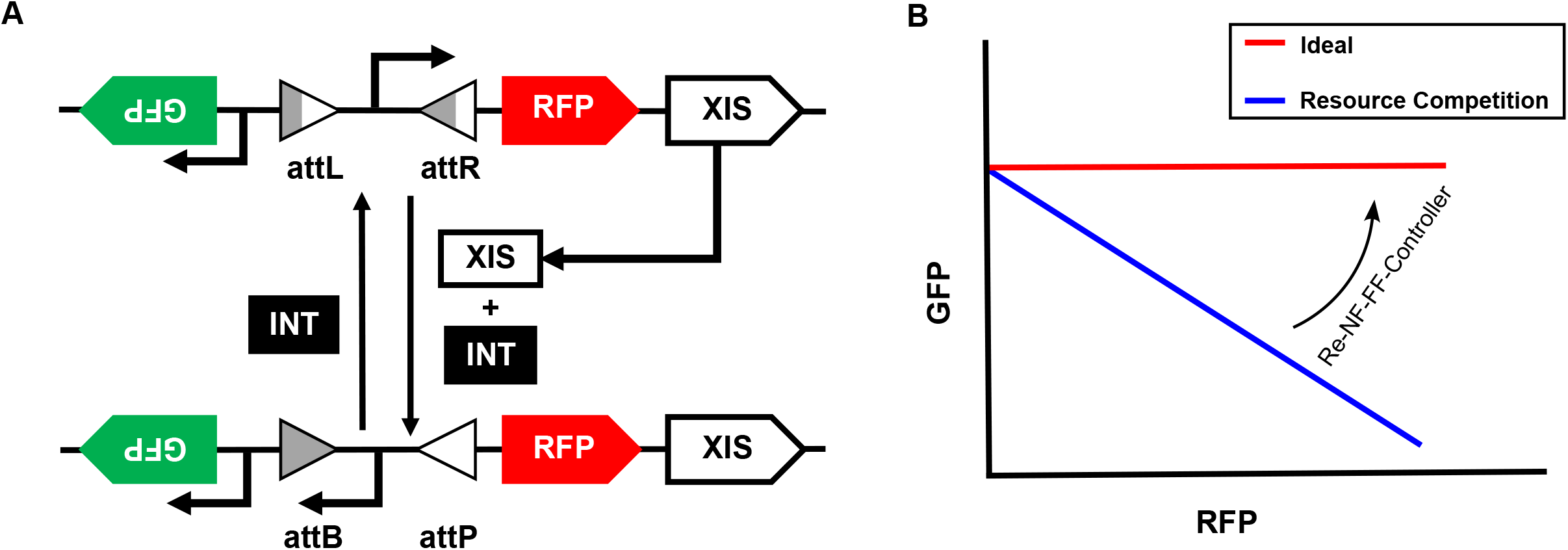
Design of a recombinase-based controller. (A) Schematic of the synthetic gene circuit, where GFP is driven by a constitutive promoter and RFP is controlled by an inducible promoter. The orientation of the inducible promoter is regulated by recombinase elements, integrase and excisionase. Excisionase, co-expressed bicistronically with RFP, forms a complex with constitutively expressed integrase to flip the promoter by converting attL and attR sites into attB and attP sites, respectively. Integrase alone reverses this process, restoring the original promoter orientation. Re-NF-Controller and Re-NF-FF-Controller are distinguished by whether the flipped promoter drives GFP expression, with only the latter inducing GFP upon promoter inversion. (B) The extent of resource coupling is assessed by examining the relative expression of RFP and GFP. A negative correlation (blue line) indicates resource competition-driven coupling, whereas a flat relationship (red line) represents an ideal case where the controller effectively eliminates resource competition.

To address this issue, we redesigned the RFP expressing promoter by flanking it with recombinase attachment sites, specifically attL (attachment Left) and attR (attachment Right). In addition, we introduced both integrase and excisionase genes. While the former is constitutively expressed, the latter is co-expressed with RFP in a bicistronic manner. This setup enables a negative feedback mechanism: Upon induction, RFP and excisionase are co-expressed due to the bicistronic design. The excisionase then forms a complex with integrase, reversing the promoter’s direction through cleavage, reversion, and re-ligation of the attachment sites, converting attL and attR into attB (Attachment Bacteria) and attP (Attachment Phage) (30, 31), as shown in **Fig.1A**. As a result, the fraction of promoters oriented to drive RFP expression is reduced, automatically limiting RFP and excisionase’s expression as resource competition intensifies. This process reverses as excisionase levels decline and the excisionase–integrase complex diminishes, allowing free integrase to catalyze the conversion of the attB and attP (BP) sites back to the attL and attR (LR) sites (26). This control strategy is referred to as the Recombinase-based Negative Feedback Controller (**Re-NF-Controller**).

Additionally, placing GFP downstream of the flipped promoter (flanked by attB/attP) enables its transcription to be enhanced by a dual-promoter effect (32, 33), compensating for the reduced expression caused by resource competition. This approach naturally establishes a feedforward mechanism using the flipped promoter without introducing additional genes that could further burden the system. By seamlessly integrating both feedback and feedforward regulation through a single promoter flip (referred to as **Re-NF-FF-Controller** thereafter), we anticipate that this new control strategy will effectively mitigate resource competition while offering a high level of tunability.

### Evaluation of Promoter Flipping Efficiency by Integrase and Excisionase

We first constructed several gene circuits to evaluate the promoter flipping efficiency by the recombinase enzymes. Here, we choose bacteriophage Bxb1 for our design, which is a temperate phage of *Mycobacterium smegmatis* and represents the best-characterized serine integrase – excisionase system (27, 35, 36). The components required for serine-mediated recombination are relatively simple, involving only the integrase and short binding sites (∼50 bp) (34). To assess Bxb1 integrase activity in our system, we constructed two circuits, C5 and C6 (**Fig. 2A**). Circuit C6 is designed to switch from GFP to RFP expression upon induction through the promoter flipping mediated by integrase, which is absent in the reference circuit C5. GFP is placed downstream of the Pbad promoter, flanked by attB and attP sites, which serve as recognition sites for integrase-mediated promoter flipping. In the presence of the inducer L-(+)-arabinose (L-ara), the transcription factor AraC activates the Pbad promoter, driving GFP expression. However, in circuit C6, integrase is also expressed, leading to promoter inversion and subsequent RFP expression. Induction experiments showed strong GFP expression in C5, whereas C6 exhibited a marked increase in RFP expression, confirming the integrase-mediated promoter reversion, as demonstrated by both fluorescence microscopy (**Fig. 2B**) and flow cytometry (**Fig. 2C**). This confirms that integrase can reliably flip the promoter flanked by BP sites to LR sites through site-specific recombination.

**Figure 2.**
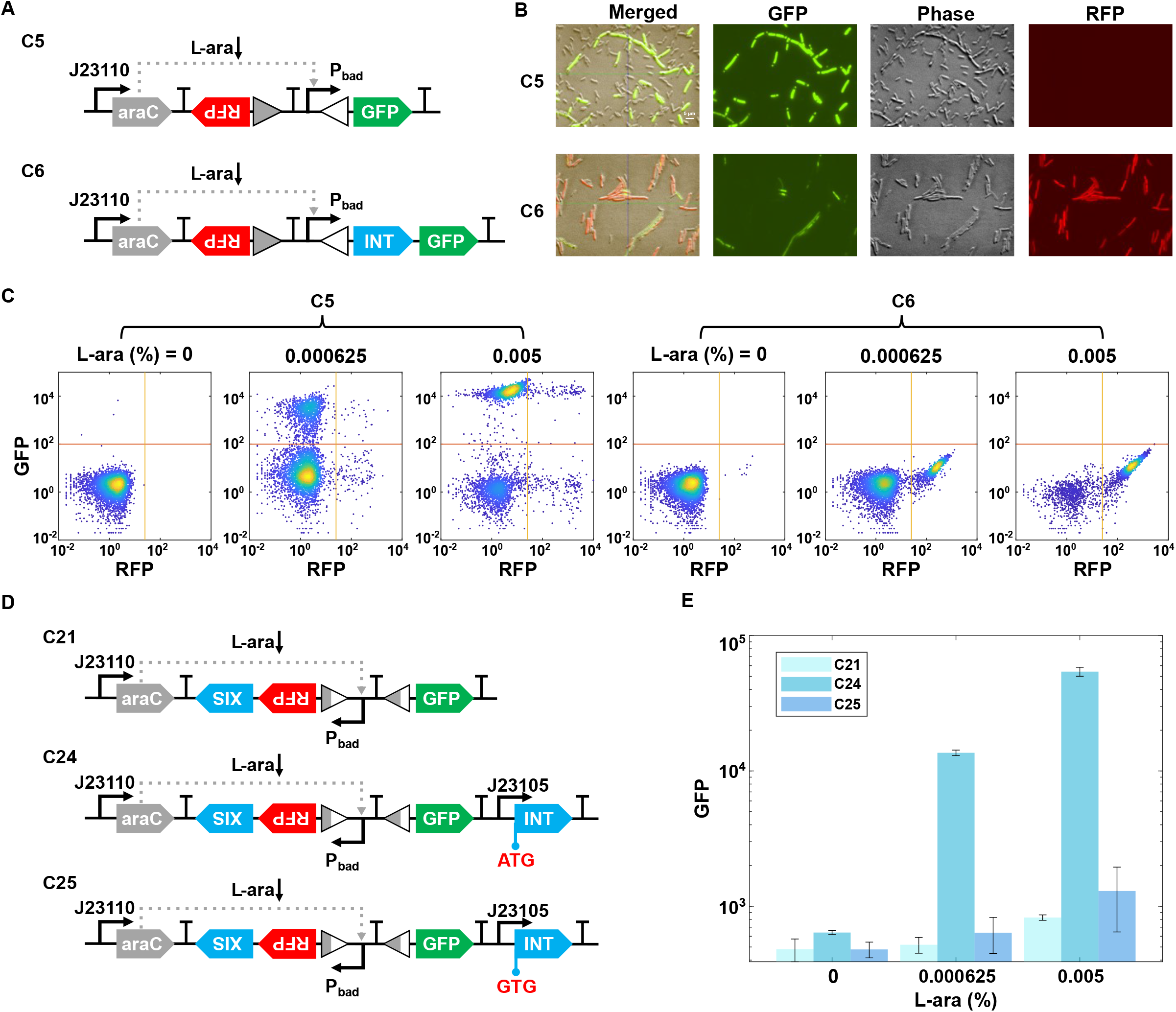
Evaluation of the flipping efficiency by the recombinase enzymes. (A) Schematic of genetic circuits C5 and C6 for testing integrase activity. Circuit C6 switches from GFP to RFP expression upon induction via promoter flipping by integrase, which is absent in the reference circuit C5. (B) Fluorescence microscopy images showing flipping efficiency. Representative results from three replicates after overnight culture with 5 × 10^−3^% L-ara concentration are shown. (C) Flow cytometry data show cell state transitions in C5 and C6 with increasing concentration of L-ara: 0%, 6.25 × 10^−4^%, and 5 × 10^−3^%. Representative results from three replicates are shown. (D) Schematic of genetic circuits C21, C24, and C25 for testing recombinase activity, with integrase constitutively expressed. Circuit C21 is the reference without recombinase. Circuit C25 uses a weaker start codon for integrase compared to circuit C24. (E) GFP expression levels in circuits C21, C24, and C25 under varying L-ara concentrations (0%, 6.25 × 10^−4^%, and 5 × 10^−3^%). Data are shown as means ± s.d. (n=3).

The conversion of LR sites back to BP sites is facilitated by excisionase, which forms a complex with integrase to specifically recognize and mediate site-specific recombination. Bxb1 gp47 was used in our design as a Bxb1-encoded excisionase that has been shown to control recombination directionality in vitro with high efficiency (27, 36). To examine the efficiency of the excisionase-directed recombination, the Pbad promoter was designed to be flanked by LR sites, driving RFP expression (**Fig. 2D**). In reference circuit C21, excisionase was co-expressed with RFP in a bicistronic arrangement upon L-ara induction. However, without integrase, the directionality of the Pbad promoter remains unchanged. In circuit C24, constitutive integrase expression allows excisionase to form a complex with it, facilitating the conversion of LR sites back to BP sites. This recombination event reorients the Pbad promoter, leading to GFP expression instead of RFP. As shown in **Fig. 2E**, upon induction, GFP levels increased significantly in circuit C24 compared with circuit C21. To further assess the role of integrase, we constructed circuit C25, which contains a weaker start codon for integrase than that in circuit C24 **(Fig. 2E)**. The results show that circuit C25 exhibited no change in GFP expression upon L-ara induction compared to circuit C24, emphasizing the necessity of fine-tuning recombinase levels to achieve optimal regulatory control.

### Recombinase-Based Negative Feedback Reduces Resource Coupling

To assess the resource decoupling capability of the Re-NF-Controller, we designed and constructed two circuits: C44 and C52 (**Fig. 3A**). In both circuits, GFP is expressed under a constitutive promoter, while RFP is regulated by the Pbad promoter. In circuit C44, negative feedback is implemented by co-expressing excisionase with the RFP module, along with a constitutively expressed integrase. Together, these enzymes can flip the orientation of the Pbad promoter, modulating RFP expression. In contrast, circuit C52 serves as reference control, where a mutation in the dinucleotide sequence of the attR site prevents recombinase-mediated flipping of the Pbad promoter, thereby eliminating the feedback mechanism. The central dinucleotide of attL and attR functions as a proofreading element during recombination and is the key determinant of substrate identity (37). Based on this, the specific dinucleotide sequence TG/AC in the attR sites was mutated to AC/TG. While recombinase can still recognize and excise both dinucleotides at the attR sites, the excised dinucleotides at the attL sites (CA/GT) can only be re-ligated with their complement pair TG/AC, but not AC/TG, at the attR sites to form the BP sites. In other words, the mutation disrupts the promoter reversion process in circuit C52 by creating a mismatch in the central dinucleotide sequences at the LR sites. As a result, the DNA fragments are unable to recombine into the BP configuration and instead re-ligate back to their original arrangement (37).

**Figure 3.**
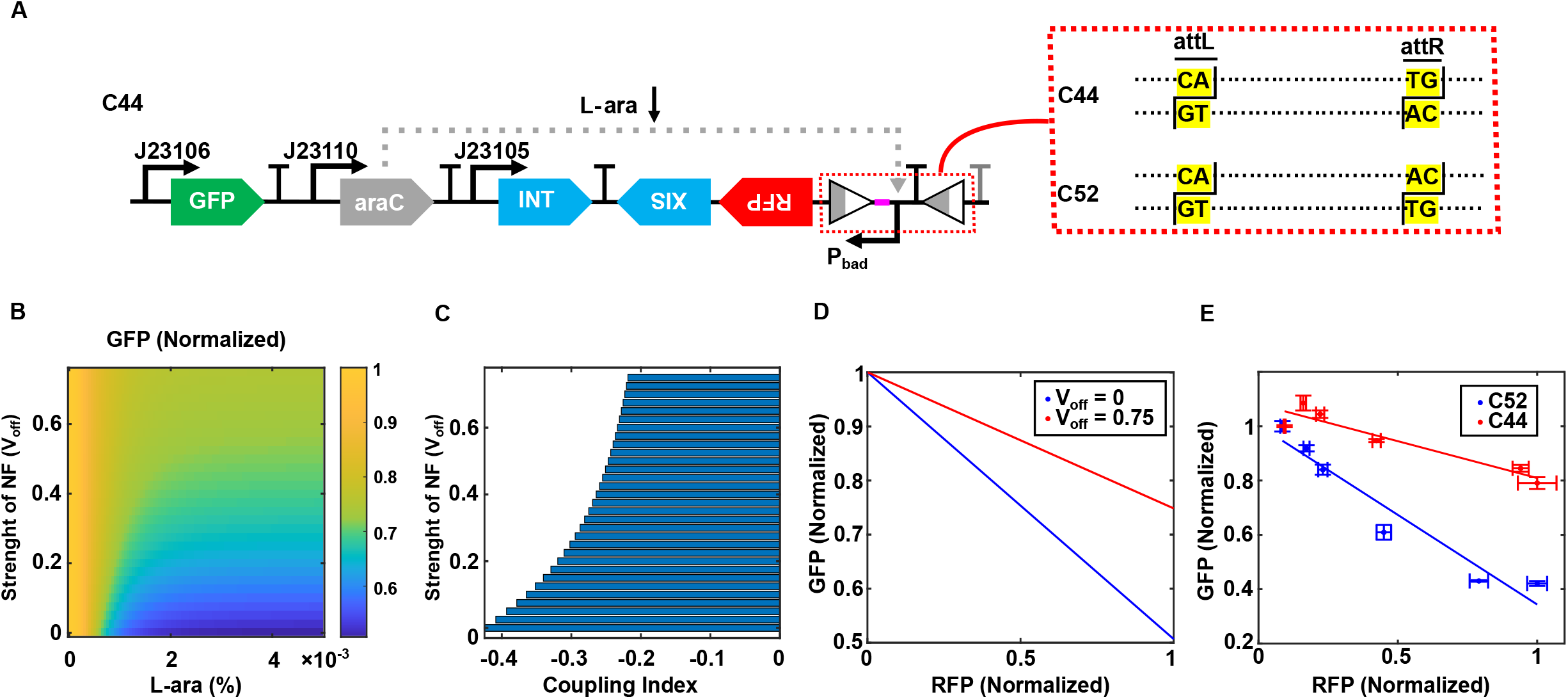
Recombinase-based negative feedback reduces resource competition. (A) Schematic representation of the open-loop circuit C52 and the genetic circuit C44 incorporating the Re-NF-Controller. C52 is derived from C44 by mutating the dinucleotide sequence at the attR site, preventing promoter flipping while maintaining the overall circuit topology. Spacer sequence (pink part) was added between attL and promoter Pbad to avoid interference. (B) Simulated steady-state GFP expression levels as a function of the inducer concentration and negative feedback strength (v_off_) in the Re-NF-Controller. (C) Reduction of the coupling index between GFP and RFP as the negative feedback strength increases in the Re-NF-Controller. (D) Comparison of the GFP-RFP correlation between the open-loop system (v_off_=0) and the system with Re-NF-Controller (v_off_=0.75). (E) Correlations between GFP and RFP expression levels in circuits C52 and C44 across different L-ara concentrations: 0%, 1.25 × 10^−4^%, 2.5 × 10^−4^%, 6.25 × 10^−4^%, 2.5 × 10^−3^%, and 5 × 10^−3^%. Solid lines represent linear fits to experimental data, while data points and error bars indicate mean ± s.d. (n=3).

To investigate how negative feedback mitigates resource-mediated coupling, we developed a mathematical model (see Methods and Supplementary Note for details). We conducted simulations to examine how GFP levels depend on both the strength of negative feedback in the Re-NF-Controller (*V*_*off*_) and the inducer dose for the RFP module (L-ara). As shown in **Fig. 3B**, in the absence of feedback (*V*_*off*_ = 0), corresponding to the open loop system, GFP levels decrease sharply with increasing L-ara, indicating significant resource competition. However, as the negative feedback strength (*V*_*off*_) increases, the dependence of GFP on L-ara weakens, reflecting reduced resource competition. To quantify this effect, we defined a coupling index as the mean change in GFP levels relative to 1. A coupling index of 0 indicates perfect decoupling, while a negative value suggests negative coupling. **Fig. 3C** shows that the coupling index is negative and increases with feedback strength until reaching a plateau. This suggests that negative feedback can effectively reduce the dependence of GFP expression on RFP expression within a certain limit. **Fig. 3D** further illustrates the relationship between GFP and RFP levels at the optimal decoupling condition. To experimentally validate this, we measured GFP and RFP expression after overnight induction of two circuits with varying L-ara concentrations. As shown in **Fig. 3E**, the GFP expression in C44 exhibited a weaker dependence on RFP levels compared to circuit C52, consistent with our modeling results. To evaluate the performance of the Re-NF-Controller under heightened resource competition, we constructed circuits C44 and C52 on a high-copy plasmid, designated C44-H and C52-H, respectively. The data show that its resource decoupling effect is diminished, revealing a limitation of this controller (**Supplementary Figure S1**). To further validate this, we also constructed another reference circuit, C42, lacking recombinase genes, with RFP expressed under the Pbad promoter and GFP driven by a constitutive promoter (**Supplementary Figure S2A**). While the GFP level as a function of RFP exhibits a flatter slope in circuit C44 compared to circuit C42 (**Supplementary Figure S2B**), the decoupling efficiency of circuit C44 compared to C42 is not as pronounced as that observed when comparing C44 to circuit C52. This is likely due to the resource consumption by controller genes, such as integrase and excisionase, in circuits C44 and C52. To assess the cellular burden imposed by the controller genes, we measured the growth rates of strains containing circuits C42 and C52 with the same gene arrangement (**Supplementary Figure S3A**). As shown in **Supplementary Figure S3B**, cells harboring C52 exhibited reduced growth compared to the C42 system, suggesting that resource consumption by integrase and excisionase imposes an unavoidable burden. Overall, theoretical and experimental analyses show that the Re-NF-Controller reduces resource coupling within certain limits.

### Terminator Malfunction Causes Unanticipated Positive Correlation in Gene Expressions

We also designed and constructed an alternative Re-NF-Controller topology by arranging gene placement to evaluate its function. In the new circuits, C43, the GFP module is positioned downstream of the Pbad promoter and attR site, separated by a terminator B0015 (**Fig. 4A)**. Surprisingly, this change in gene placement results in a completely different gene expression pattern. As shown in **Fig. 4B**, GFP expression in circuit C43 now exhibits a positive correlation with RFP level, in contrast to the negative correlation observed previously, suggesting a change in the dynamics of resource allocation toward the GFP module.

**Figure 4.**
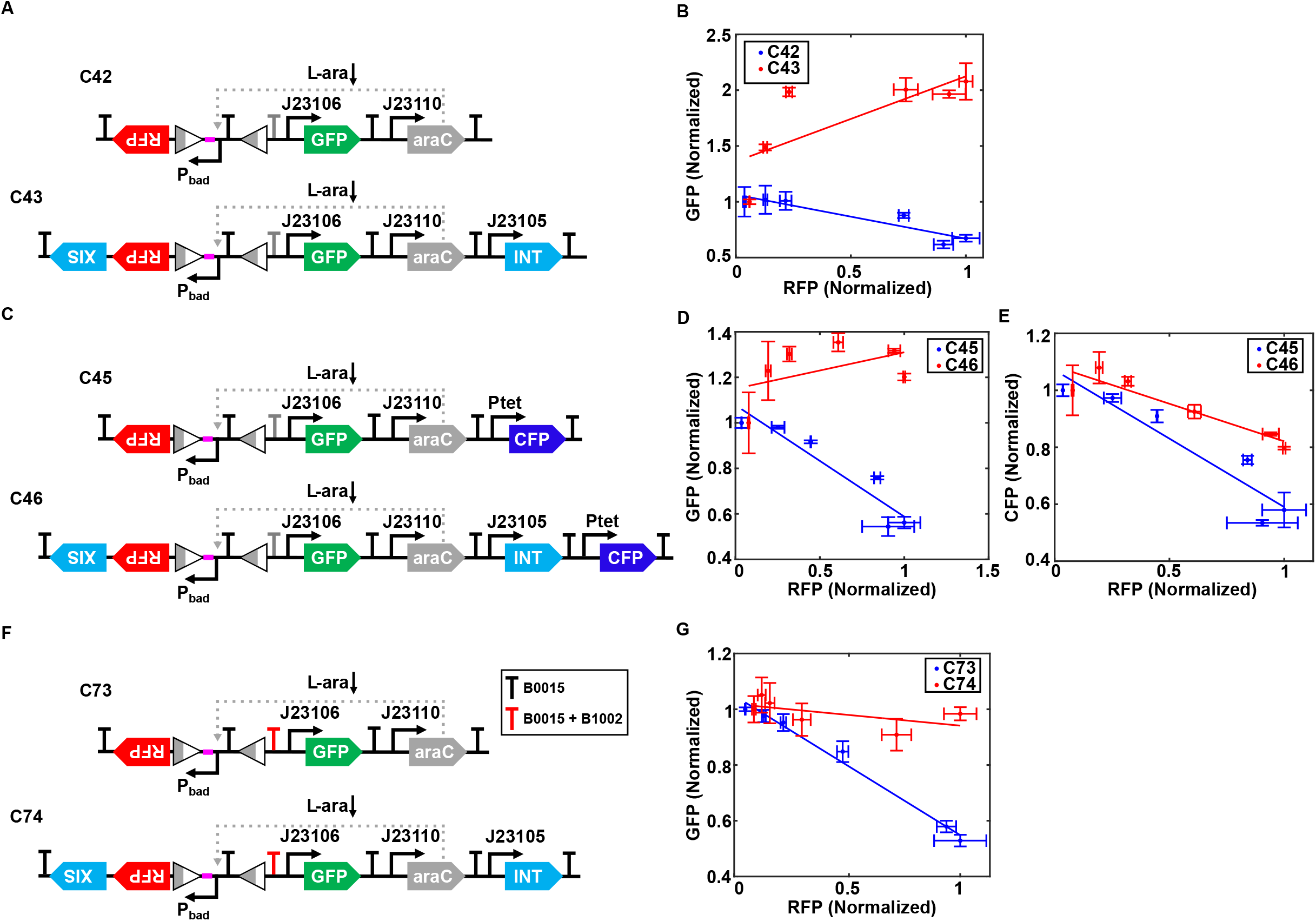
Malfunction of the terminator leads to overcorrection in resource decoupling. (A) Schematic of genetic circuit C43, an alternative Re-NF-Controller design where the GFP module is placed downstream of the Pbad promoter and attR site. Circuit C42 serves as a reference without recombinase elements. Spacer sequence (pink part) was added between attL and promoter Pbad to avoid interference. (B) Correlations between GFP and RFP expression levels in circuits C42 and C43 across different L-ara concentrations: 0%, 1.25 × 10^−4^%, 2.5 × 10^−4^%, 6.25 × 10^−4^%, 2.5 × 10^−3^%, and 5 × 10^−3^%. Solid lines represent linear fits to experimental data, while data points and error bars indicate mean ± s.d. (n=3). (C) Schematic diagram of genetic circuits C45 and C46, incorporating constitutively expressed CFP as an independent indicator to assess resource coupling in the system. Malfunctional terminators were labeled grey in all figures. (D-E) The correlation between GFP (d) or CFP (e) expression level and RFP expression level in circuits C45 and C46. The experimental conditions were the same as those described in panel (b). (F) Schematic diagram of genetic circuits C73 and C74, incorporating double terminators between the Pbad promoter and the GFP module. (G) The correlations between GFP and RFP expression levels in C73 and C74. The experimental conditions were the same as those described in panel (b).

To validate the issue of placement dependent, we introduced an additional fluorescence reporter gene, cyan fluorescent protein (CFP), into circuits C42 and C43, positioned far from the attR sites, resulting in circuits C45 and C46 (**Fig. 4C**). Experimental data show that the GFP level still increases abnormally with RFP expression in circuit C46 (**Fig. 4D**). However, the CFP level exhibits the expected negative correlation with RFP expression, with a less steep slope compared to C45, indicating the normal function of the Re-NF-Controller (**Fig. 4E**). These distinct expression patterns between CFP and GFP may strongly stem from the upstream spacer, which added in front of the Pbad promoter in circuits C42 and C43 or the loss of terminator function downstream of the attR site.

To assess whether the effect is caused by the spacers, we constructed circuits C34 and C35 with the spacers removed. A similar induction pattern was observed (**Supplementary Figure S4A**), except for differences in the relative expression of RFP and GFP. Upon spacer removal, the differential RFP expression between the unregulated and recombinase-regulated circuits is enhanced (**Supplementary Figure S4B**) and GFP expression is reduced (**Supplementary Figure S4C**). We also tested whether adding a fast degradation tag to excisionase would resolve the issue (**Supplementary Figure S5A**). However, we found that resource coupling becomes unaffected **(Supplementary Figure S5B-C)**, These data suggest that while the spacer helps mitigate resource competition but is not responsible for the abnormal relationship between GFP and RFP. We further tested the functionality of terminator B0015 by constructing two simple circuits (C65 and C66) with GFP placed downstream of the flipped Pbad promoter, either with or without the terminator, respectively (**Supplementary Figure S6A**). As shown in **Supplementary Figure S6B**, no difference was observed in the induced GFP expression pattern as a function of L-ara dose between two circuits, suggesting that terminator B0015 had lost its function after the attR site.

To resolve this issue, we inserted an additional short terminator, B1002, downstream of the existing B0015 terminator, generating circuit C70 (**Supplementary Figure S6A**). This dual-terminator design prevented GFP expression from the flipped Pbad promoter, as shown in **Supplementary Figure S6B**. We then redesigned circuits C42 and C43 by incorporating the dual terminator, resulting in circuits C73 and C74 (**Fig. 4F**). In these redesigned circuits, we observed the expected negative correlation between GFP and RFP expression, with a reduced slope in C73 (**Fig. 4G**), as opposed to the positive correlation seen previously. **Supplementary Figure S7** shows the coupling index, illustrating the enhanced resource decoupling of circuits C46 and C74 compared to their reference circuits.

### Additional Feedforward Control Enhances Mitigation of Resource Competition

The above observation was unexpected but can be leveraged for further minimizing resource coupling. By removing the B0015 terminator in C43, we constructed circuit C53 (**Fig. 5A**). In this new design, the promoter reversion induced by the integrase-excisionase complex not only limits RFP expression through a negative feedback mechanism but also compensates GFP expression via a feedforward control mechanism. We also constructed a reference circuit C55, which lacks the promoter-inverting function due to mutated dinucleotides in the attR sites. By combining the negative feedback and feedforward mechanisms, we hypothesize that this new design, “**Re-NF-FF-Controller**”, can enhance the efficiency of resource competition mitigation. By fine-tuning the recombinase elements, we can correct the overcompensation for the GFP module, ensuring that its expression is largely independent of RFP expression, exhibiting neither positive nor negative correlation.

**Figure 5.**
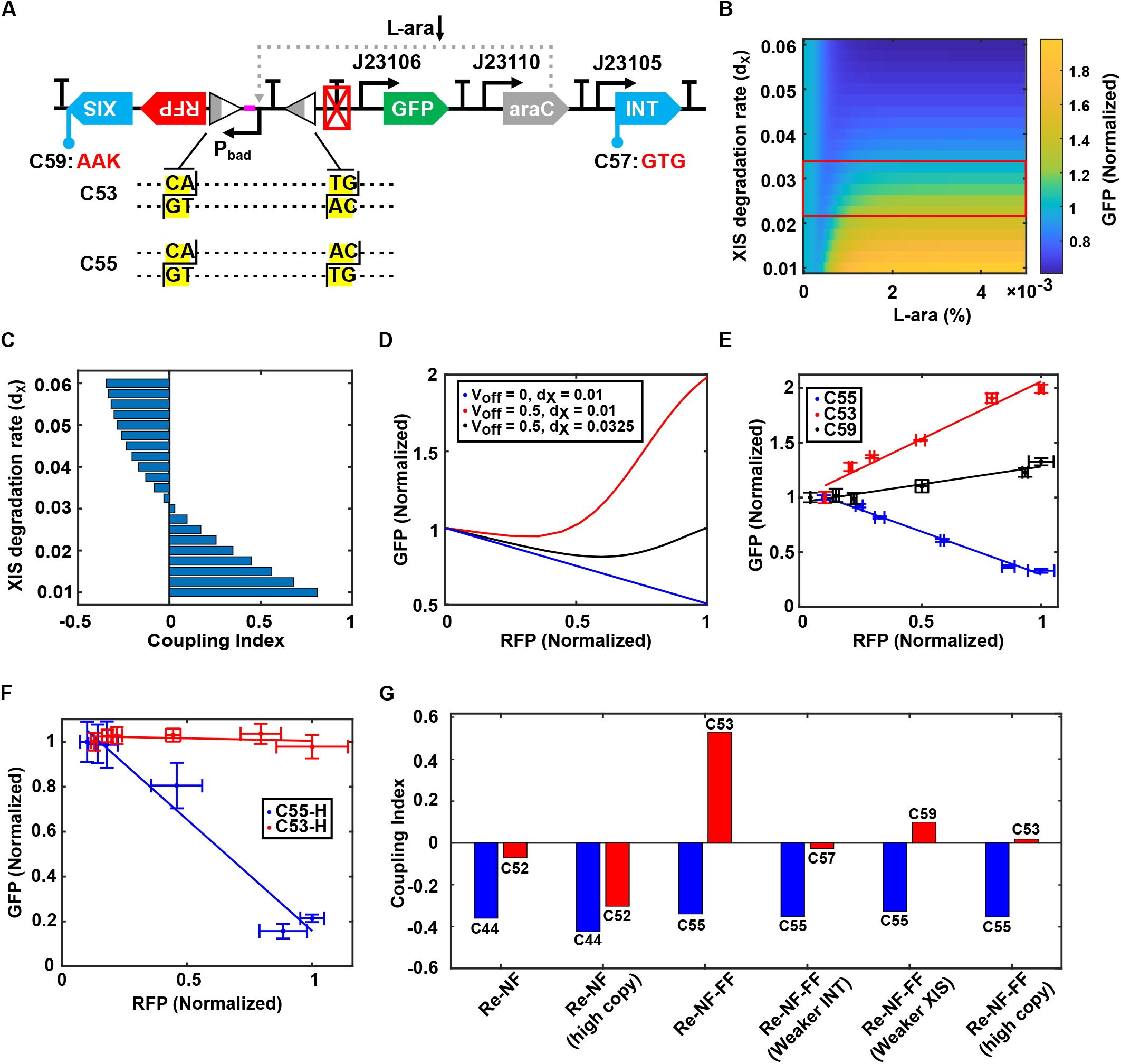
Additional Feedforward Control Significantly Enhances Resource Decoupling. (A) Schematic diagram of genetic circuits C53, C55, C57, and C59. Circuit C53 incorporates the Re-NF-FF-Controller, while C55 represents its corresponding open-loop system. Circuit C57 features a modified integrase start codon (ATG to GTG), resulting in reduced integrase expression. Circuit C59 includes a degradation tag (AAK) added to excisionase, enabling rapid degradation of excisionase. (B) Simulated steady-state GFP expression levels as a function of inducer concentration and the degradation rate of excisionase (dx). The boxed region highlights the effective resource decoupling achieved by the Re-NF-FF-Controller. (C) The coupling index between GFP and RFP as a function of the degradation rate of excisionase (d_x_). (D) The optimal resource decoupling effect achieved by tuning the degradation rate of excisionase to correct the overcompensation (red line). The open-loop system (blue line) is shown as a reference. (E) Correlations between GFP and RFP expression levels in circuits C53, C55, and C59 across different L-ara concentrations: 0%, 1.25 × 10^−4^%, 2.5 × 10^−4^%, 6.25 × 10^−4^%, 2.5 × 10^−3^%, and 5 × 10^−3^%. (F) Correlation between GFP and RFP expression in high-copy circuits C53-H and C55-H. Induction conditions are as described in panel (E). Solid lines represent linear fits to experimental data; data points and error bars indicate mean ± s.d. (n = 3). (G) Coupling index quantifying the resource decoupling performance of Re-NF and Re-NF-FF controllers under various conditions.

We developed a mathematical model to demonstrate the tunability of the Re-NF-FF-Controller in regulating resource competition. The GFP level, mapped against the inducer dose of the RFP module (L-ara) and the degradation rate of excisionase (dx), reveals that optimal resource decoupling can be achieved through fine-tuning of dx (red boxed area, **Fig. 5B**). If the degradation rate of excisionase is too slow, the feedforward mechanism becomes excessively strong, leading to overcompensation and a positive coupling index (**Fig. 5C**). Conversely, if the degradation rate is too fast, the negative feedback loop becomes too weak to effectively reduce resource coupling, resulting in a system that still exhibits a negative coupling index (**Fig. 5C**). By optimizing excisionase level, the Re-NF-FF-Controller can maintain relatively stable GFP expression, minimizing its dependence on RFP expression (**Fig. 5D**). A similar strategy can be applied by fine-tuning integrase expression levels. As shown in **Supplementary Figure S8A-C**, optimal integrase expression rate minimizes GFP perturbation and the coupling index. However, excessively high or low integrase expression leads to strong positive or negative correlations, respectively. These simulation results underscore the importance of maintaining balanced recombinase expression for the Re-NF-FF-Controller to effectively regulate resource competition.

To experimentally validate the simulation results, we constructed circuit C59 by introducing a degradation tag to excisionase (**Fig. 5A**). Analyzing gene expression patterns across these systems revealed distinct correlations between GFP and RFP levels. The reference circuit C55 exhibited a strong negative correlation due to resource competition, while circuit C53 showed a strong positive correlation caused by overcompensation. In contrast, circuit C59 demonstrated a more balanced expression pattern, with a relatively flattened curve compared to both C53 and C55 (**Fig. 5E**). To validate promoter flipping and its timing, we performed PCR amplification following induction of circuits C53 and C55 at different time points (see Methods). As shown in **Supplementary Figure S9**, the attBP band appeared at 9 hours post-induction in C53 but was absent in the reference circuit C55, indicating that promoter flipping occurs between 6 and 9 hours after induction.

To evaluate the performance of the Re-NF-FF-Controller under heightened resource competition, we constructed high-copy versions of circuits C53 and C55, termed C53-H and C55-H respectively. As shown in **Fig. 5F**, the reference circuit C55-H exhibited a very strong negative correlation between GFP and RFP expression, indicating increased resource competition. In contrast, C53-H showed minimal dependence of GFP on RFP levels, demonstrating effective decoupling. These results suggest that fine-tuning excisionase levels in response to elevated resource limitation can improve the decoupling performance of the Re-NF-FF-Controller.

To test whether modulating integrase level could yield a similar tuning effect, we constructed circuit C57 by weakening integrase expression through a start codon mutation from ATG to GTG (**Fig. 5A**). As a result, GFP expression in C57 showed the weakest dependence on RFP compared to C53 and C55, indicating enhanced resource decoupling (**Supplementary Figure S8D**). The overall resource-decoupling performance of the Re-NF and Re-NF-FF controllers across different conditions is compared by the coupling index (CI). As shown in **Fig. 5G,** while the Re-NF controller exhibits strong decoupling, its performance declines under high resource competition with increased plasmid copy number. In contrast, the Re-NF-FF-controller suffers from overcompensation but can be significantly improved through fine-tuning of integrase or excisionase expression. Overall, compared to the Re-NF-controller, the Re-NF-FF-controller offers greater resilience in resource-limited environments and features a tunable design for optimized performance. Taken together, the Re-NF-FF-Controller integrates both negative feedback and feedforward mechanisms to achieve tunable resource distribution, effectively reducing resource competition in synthetic gene circuits.

## Discussion

Resource competition adds another layer of complexity to engineering robust synthetic circuits. In this study, we observed substantial resource competition in open-loop systems, consistent with previous findings (6, 8). To address this, we introduced a novel control strategy that integrates recombinase elements to restore the modularity of gene modules through a combination of negative feedback and feedforward regulation. Our results strongly suggest that combining feedforward with negative feedback effectively overcomes the limitations of negative feedback alone.

Negative feedback is a fundamental regulatory motif in biological networks, known for its role in maintaining homeostasis and minimizing variability (28, 38–40). Although this strategy can mitigate resource competition and reduce coupling, its capacity is inherently limited by controller saturation, where the regulatory elements (e.g., repressors or activators) reach a maximum effect and can no longer proportionally respond to increasing perturbations, thus failing to achieve full decoupling. This limitation was also evident in our recombinase-based negative feedback systems, highlighting the need for a more efficient regulatory topology. The combination of multiple regulatory mechanisms to achieve desired outcomes is a common strategy in synthetic biology. For example, negative feedback is coupled with positive feedback to achieve robust bistability, oscillation, and adaptation (41, 42). By coupling a negative feedback loop with incoherent feedforward control, Yang et al. designed a circuit called “Equalizer” to buffer plasmid copy number variation (43).

However, combining these two regulatory mechanisms to control resource coupling in synthetic gene circuits presents challenges. First, more complex architecture inevitably introduces intricate interactions in circuit design and engineering. For instance, single-promoter circuits tend to be more robust than multi-promoter circuits (44). Therefore, without compromising circuit function, a simple yet effective design can outperform a more complex one. In our case, using a single promoter (Pbad) instead of multiple promoters for each regulatory module simplifies circuit construction. This minimalistic design also facilitates integration into larger genetic networks by reducing the likelihood of unpredictable interactions that often arise from circuit complexity. Additionally, the use of more gene parts to construct complex structures to implement multiple control mechanisms consumes considerable cellular resources, imposing a significant burden on the host cells. Our design only requires the reversion of a single promoter by recombinase to achieve both regulatory mechanisms, avoiding the need for additional genes/promoters that would demand extra cellular resources. This minimizes the metabolic burden on the host cell. Furthermore, the simplicity of the system does not compromise its flexibility, as the desired output can be fine-tuned by adjusting the levels of integrase and excisionase.

Like all control systems that incorporate negative feedback, the expression level of the regulated gene will be lower than the open-loop system (11, 16, 24, 28). This challenge can be addressed by increasing the translational strength of regulated genes. Since recombinase expression imposes a cellular burden, exogenous expression or chromosomal integration could further improve efficiency and robustness. Nonetheless, the proposed recombinase-based resource-regulating strategy holds great potential for future applications. Zhao et al. proposed a binary ripple counter using integrase and its directionality factor to toggle states for event recording, which could be extended for recording large numbers of events using multiple orthogonal recombinases (45). More than 25 putative large serine recombinases have been identified (46–48). For each recombinase, up to six orthogonal recombination site pairs can be engineered by introducing distinct non-symmetric 2-bp central overlap sequences. As a result, each attachment site recombines exclusively with its corresponding partner (49). A comprehensive evaluation of these recombinases and their corresponding sites could enable the construction of scalable, resource-aware genetic circuits. Furthermore, recombinase elements have been successfully applied in species like mammals and yeast (50, 51), broadening the potential for recombinant-based regulation in diverse systems.

## Data availability

All data produced or analyzed for this study are included in the article and its Supplementary Information files. The simulation code is shared at GitHub (https://github.com/TianLab-ASU/Re-NF-FF-Controller).

## Acknowledgments

This work was supported by grants from the US National Institutes of Health (R35GM142896 to X.-J.T.).

## Author contributions

X.-J.T. conceived the study. X.-J.T., R.Z. and RX.Z. designed the study. R.Z. and RX.Z. performed experiments. X.-J.T. and RX.Z. performed theoretical studies. X.-J.T., R.Z. and RX.Z. analyzed the data. X.-J.T. and RX.Z. wrote the manuscript. X.-J.T., R.Z. and RX.Z. edited the manuscript.

