## Supplementary Information for "Integrating Recombinase-Based Feedback and Feedforward Control for Optimal Resource Decoupling"

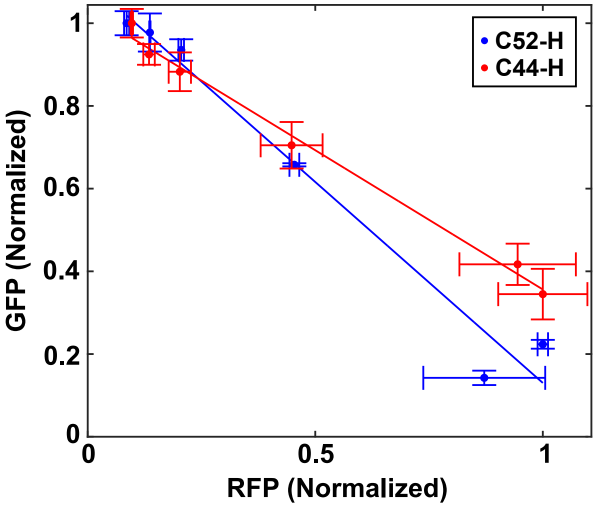


**Supplementary Fig.1. Effect of high copy plasmid on the resource decoupling efficiency of Re-NF-Controller.** Comparison of the correlations between GFP and RFP expression levels in circuits C44 and C52 with high plasmid copy numbers (C44-H and C52-H) across different L-ara concentrations: 0%, $1.25 \times 10^{-4}$%, $2.5 \times 10^{-4}$%, $6.25 \times 10^{-4}$%, $2.5 \times 10^{-3}$%, and $5 \times 10^{-3}$%. Solid lines represent fitted experimental data, while data points and error bars indicate mean ± s.d. (n=3).

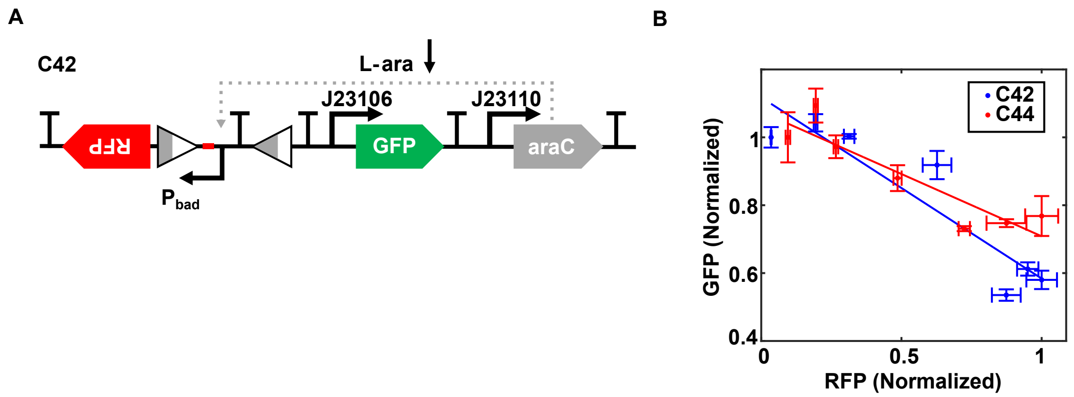


**Supplementary Fig. 2. Recombinase-based negative feedback reduces resource competition in gene circuits, compared to the open-loop system without recombinase.**

(A) Schematic diagram of reference circuit C42, which is an open-loop system without recombinase.

(B) Comparison of the correlations between GFP and RFP expression levels in circuits C42 and C44 across different L-ara concentrations: 0%, $1.25 \times 10^{-4}$%, $2.5 \times 10^{-4}$%, $6.25 \times 10^{-4}$%, $2.5 \times 10^{-3}$%, and $5 \times 10^{-3}$%, $8.3 \times 10^{-3}$%. Solid lines represent fitted experimental data, while data points and error bars indicate mean ± s.d. (n=3).

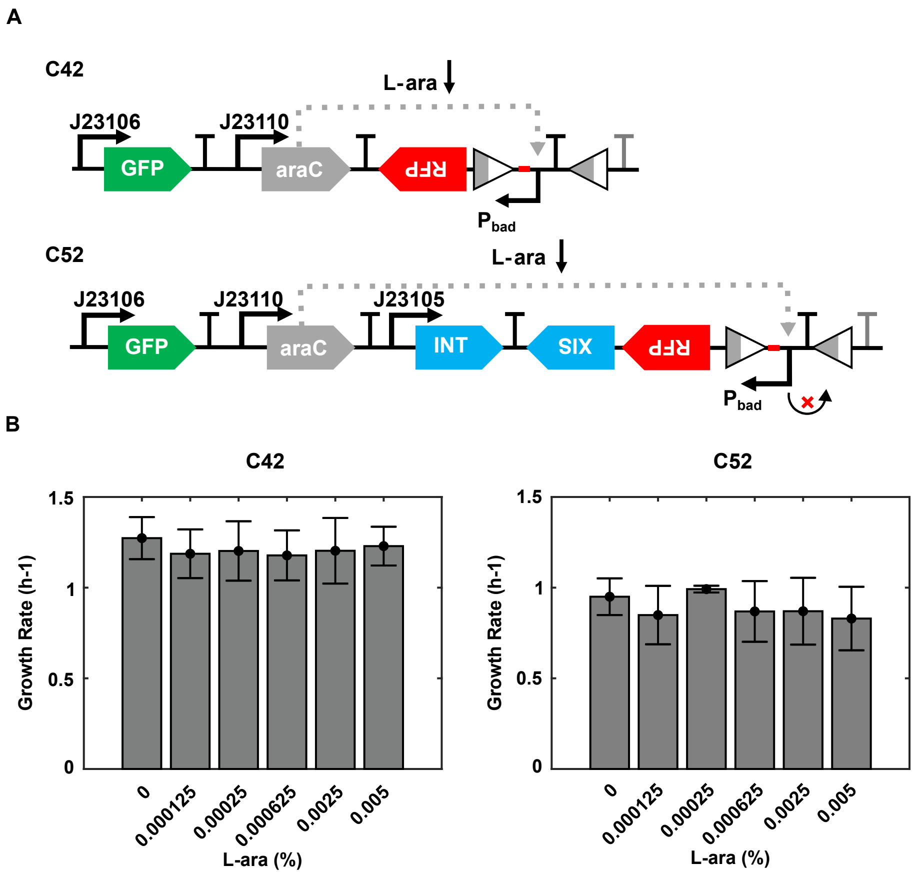


**Supplementary Fig. 3. Assessment of controller gene burden as indicated by the exponential growth rate of host cells.**

(A) Schematic diagram of gene circuits. The topology of C42 was rearranged to match that of C52, allowing the assessment of the burden imposed solely by the controller genes.

(B) The exponential growth rates of C42 and C52 were dynamically measured across varying L-ara concentrations: 0%, $1.25 \times 10^{-4}$%, $2.5 \times 10^{-4}$%, $6.25 \times 10^{-4}$%, $2.5 \times 10^{-3}$%, and $5 \times 10^{-3}$%. The experimental data and error bars represent means ± s.d., n=3.

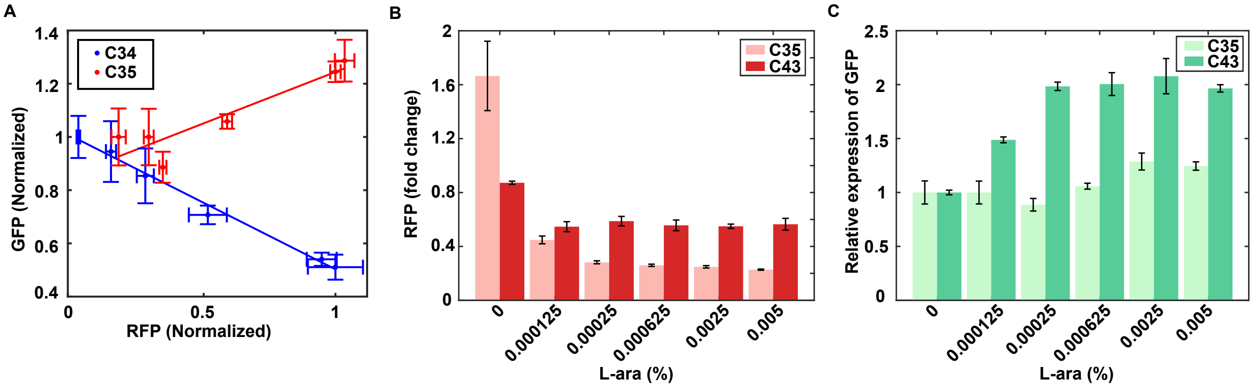


**Supplementary Fig. 4. Evaluation of the impact of spacer insertion on gene expression in recombinase-modified gene circuit.**

(A) The correlation between GFP and RFP expression levels in circuits C34 and C35, which are similar to C42 and C43, but with the spacer between the Pbad promoter and attL site removed. The experiments were performed across different L-ara concentrations: 0%, $1.25 \times 10^{-4}$%, $2.5 \times 10^{-4}$%, $6.25 \times 10^{-4}$%, $2.5 \times 10^{-3}$%, and $5 \times 10^{-3}$%. Solid lines represent fitted experimental data, while data points and error bars indicate mean ± s.d. (n=3).

(B) Comparison of the fold change ($log_2$) of RFP expression between circuits without spacer (C35) and with spacer (C43), relative to their respective reference circuits C34 and C42 respectively.

(C) Comparison of the fold change of GFP expression in circuits without spacer (C35) and with spacer (C43) based on the expression in the absence of inducer.

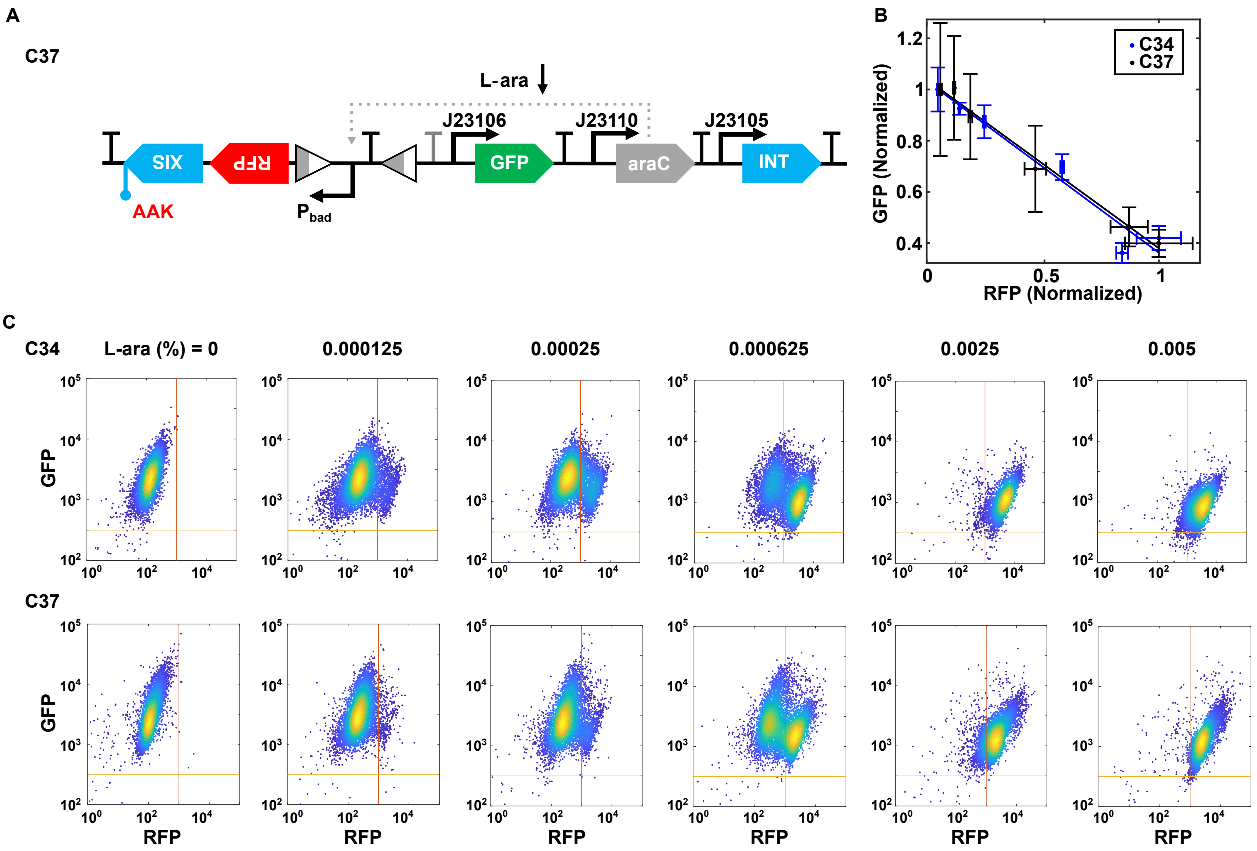


**Supplementary Fig. 5. Testing the Re-NF-Controller System with a Fast Degradation Tag on Excisionase.**

(A) Schematic of genetic circuit C37, where a fast degradation tag (AAK) is fused to excisionase for enhanced degradation.

(B) The correlation between GFP and RFP expression in circuit C37, showing no resource decoupling compared with the reference circuit C34. The experiments were performed across different L-ara concentrations: 0%, $1.25 \times 10^{-4}$%, $2.5 \times 10^{-4}$%, $6.25 \times 10^{-4}$%, $2.5 \times 10^{-3}$%, and $5 \times 10^{-3}$%. Solid lines represent fitted experimental data, while data points and error bars indicate mean ± s.d. (n=3).

(C) Flow cytometry data shows no change in GFP expression levels despite increased RFP expression upon L-ara induction in circuit C37, similar to the reference circuit C34. Representative results from three replicates are shown.

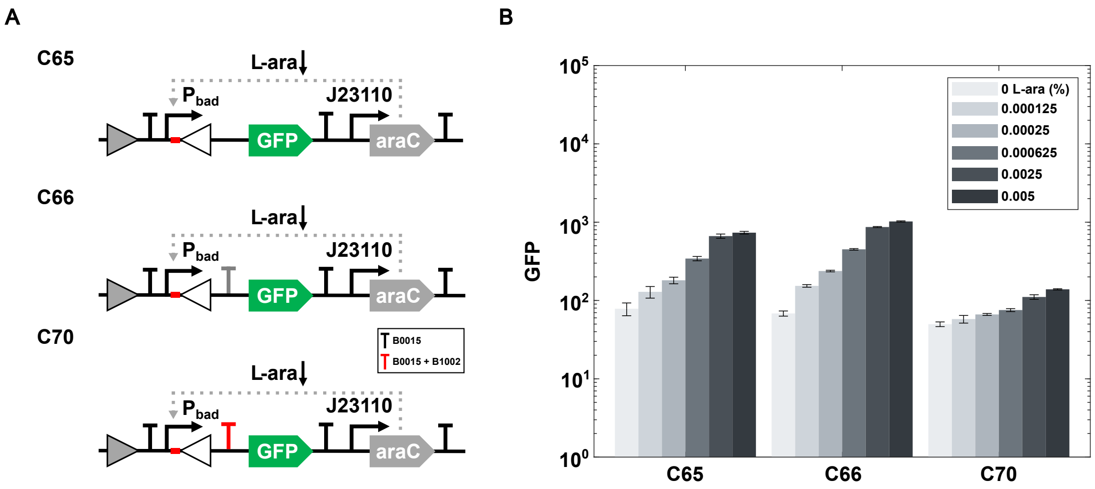


**Supplementary Fig. 6. Evaluation of terminator efficiency downstream of recombinase sites after promoter flipping.**

(A) Schematic diagram of genetic circuits: no terminator (C65), one terminator (B0015) (C66), or double terminators (C70) inserted downstream of Pbad.

(B) GFP expression upon induction with increasing L-ara concentrations (0%, $1.25 \times 10^{-4}$%, $2.5 \times 10^{-4}$%, $6.25 \times 10^{-4}$%, $2.5 \times 10^{-3}$%, and $5 \times 10^{-3}$%) measured using a plate reader. The experimental data and error bars represent means ± s.d., n=3.

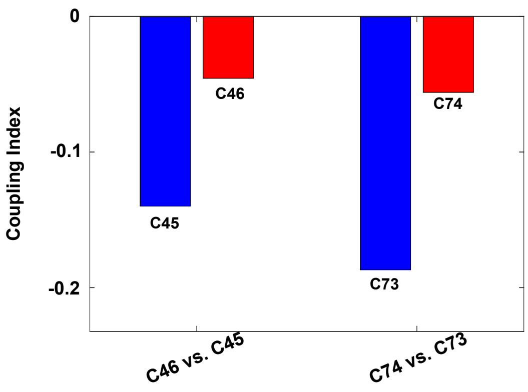


**Supplementary Fig. 7.** Coupling index shows the resource decoupling effect of the circuits C46 and C74 compared to their reference circuits.

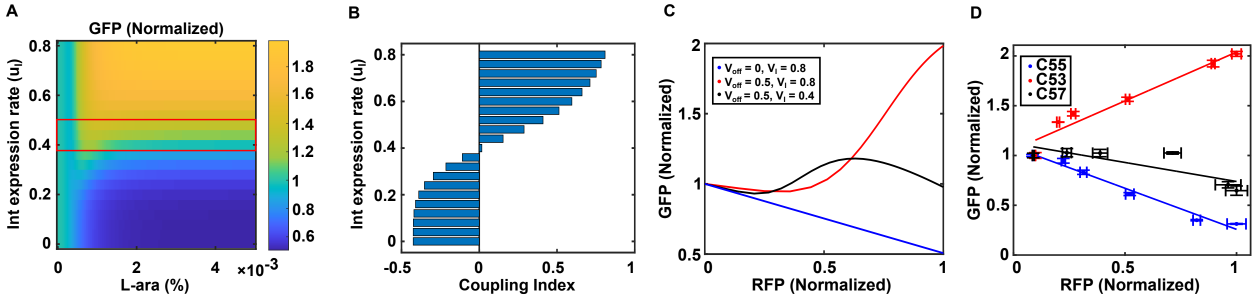


**Supplementary Fig. 8. Optimizing resource competition mitigation in the Re-NF-FF-Controller by tuning the integrase transcriptional,**

(A) Simulated steady-state GFP expression levels as a function of inducer concentration and expression rate of integrase. The boxed region highlights the effective resource decoupling achieved by the Re-NF-FF-Controller.

(B) The coupling index between GFP and RFP as a function of the expression rate of integrase.

(C) The optimal resource decoupling effect achieved by tuning the expression rate of integrase (black line) to correct the overcompensation (red line). The open-loop system (blue line) is shown as a reference.

(D) Correlations between GFP and RFP expression levels in circuits C53, C55, and C57 across different L-ara concentrations: 0%, $1.25 \times 10^{-4}$%, $2.5 \times 10^{-4}$%, $6.25 \times 10^{-4}$%, $2.5 \times 10^{-3}$%, and $5 \times 10^{-3}$%. Solid lines represent fitted experimental data, while data points and error bars indicate mean ± s.d. (n=3).

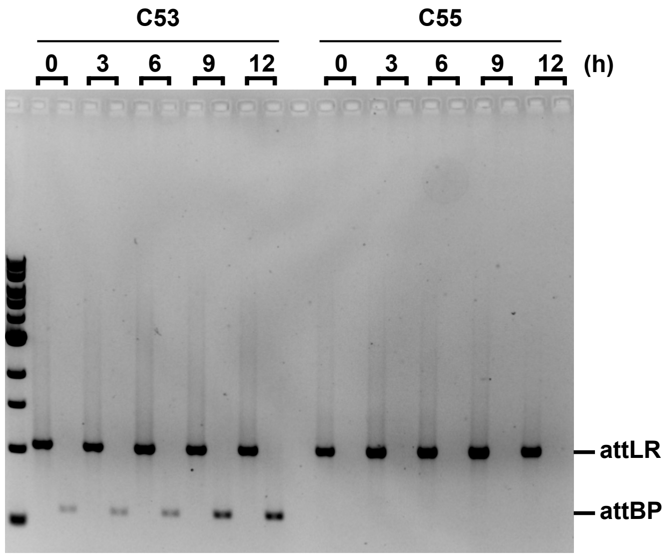


**Supplementary Fig. 9. Gel electrophoresis analysis of PCR products to determine the timing of promoter flipping in circuits C53 and C55.** The system with circuit C53 or C55 was induced with $5 \times 10^{-3}$% L-ara at different time series: 0, 3, 6, 9, 12h.

### Mathematical modeling

To simulate the effect of recombinase-based control system on controlling resource competition in cellular environment, we built a mathematical model utilizing a system of ordinary differential equations (ODE).

#### 1. Mathematical Modeling of Resource Competition in the System

Here, we present the general mathematical model for resource competition in a system, which will be used to describe and analyze system dynamics both in the absence of a controller and with the Re-NF-Controller or Re-NF-FF-Controller in subsequent sections. The dynamics of protein concentration $[P_i]$ in the system are modeled as follows:

$$\frac{d[P_i]}{dt} = v_i * \frac{CN_i * R_i/Q_i}{PF_Q} - d_i * [P_i]$$

Where $CN_i$ is gene copy number, $v_i$ is the expression rate of protein $P_i$, and $d_i$ is the degradation rate of protein $P_i$. $R_\mathrm{i}$ is the fraction of active promoters for expressing protein $P_i$. For constitutive promoters, $R_\mathrm{i} = 1$. For inducible promoters, such as Pbad in our system, $R_\mathrm{i}$ depends on the concentration of the transcription factor, following a Hill function. For example, consider the protein controlled by Pbad promoter, $\mathrm{R_i} = \frac{Sa*[AraC]^2}{Sa*[AraC]^2+1}$, where $Sa = C_{min} + (C_{max} - C_{min}) * \left(\frac{Lara^n}{Lara^n+J^n}\right)$. $Sa$ describes how the promoter activity is regulated by inducer L-ara, Cmax and Cmin are the maximum and minimum affinities of the AraC dimers to the binding sites on the promoter Pbad. The details about model construction for AraC activated system are further explained in our previous work (1).

It is noted that the effect of the resource competition is formulated with $\mathrm{PF_Q} = 1 + CN\sum\frac{R_\mathrm{i}}{Q_\mathrm{i}}$, where $Q_i$ is introduced to represent the difference in resource competitivity between genes. The derivation of the equations describing resource competition can be found in our previous work (2). Building on this modeling framework described, we developed models for three systems: the open-loop system, the system with the Re-NF-Controller, and the system with the Re-NF-FF-Controller.

#### 2. Mathematical Model for the Open-Loop System.

In the Open-Loop System, promoter flipping does not occur. We applied the general resource competition model to all proteins in the system, leading to the following set of equations:

$$\frac{d[INT]}{dt} = u_I * \frac{CN/Q_I}{PF_Q} - d_I * [INT]$$

$$\frac{d[XIS]}{dt} = u_R * \frac{CN * R_R/Q_R}{PF_Q} - d_X * [XIS]$$
$$\frac{d[AraC]}{dt} = u_A * \frac{CN/Q_A}{PF_Q} - d_A * [AraC]$$
$$\frac{d[GFP]}{dt} = u_{G1} * \frac{CN/Q_G}{PF_Q} - d_G * [GFP]$$
$$\frac{d[RFP]}{dt} = u_R * \frac{CN * R_R/Q_R}{PF_Q} - d_R * [RFP]$$

Where $PF_Q = 1 + CN * \left(\frac{1}{QI} + \frac{1}{Q_A} + \frac{1}{Q_G} + \frac{2*R_R}{Q_R}\right), R_R = \frac{Sa*[AraC]^2}{Sa*[AraC]^2+1}, Sa = C_{min} + (C_{max} - C_{min}) * \left(\frac{Lara^n}{Lara^n+J^n}\right)$.

**3. Mathematical Model for the System with Re-NF-Controller**

In the system with a Re-NF-Controller, the orientation of the Pbad promoter can be flipped by recombinase. Initially, all copies of the Pbad promoter are designed in the LR state to drive RFP expression. Upon induction, RFP and excisionase accumulate in the system, allowing excisionase to form a complex with the constitutively expressed integrase, which flips the Pbad promoter into the BP state. We introduce an equation to model the proportion of Pbad promoter copies remaining in the LR state $\phi \in [0,1]$

$$\frac{d[\phi]}{dt} = v_{\text{on}} * \frac{[INT]^4}{{K_{I1}}^4 + [INT]^4} * (1 - \phi) - v_{\text{off}} * \frac{[XIS]^4}{{K_X}^4 + [XIS]^4} \frac{[INT]^4}{{K_{I2}}^4 + [INT]^4} * \phi$$

where the first and second terms represent the promoter flipping rate between LR and BP states, with maximum rate constant of $v_{\text{on}}$ and $v_{\text{off}}$, respectively. $K_{I1,2}$ and $K_X$ are equilibrium constants that determine the integrase and excisionase concentration required to reach half-maximal flipping rate (3, 4). Since four integrase molecules are required for promoter flipping, the integrase-dependent fraction follows a fourth-order Hill function. The integrase-excisionase complex is assumed to have a 1:1 stoichiometry for catalyzing LR-to-BP recombination, leading to an excisionase-dependent fraction modeled by a Hill function with a coefficient of 4.

The equation for RFP needs to be revised accordingly

$$\frac{d[RFP]}{dt} = u_R * \frac{CN * \phi * R_R/Q_R}{PF_Q} - d_R * [RFP]$$

To simulate how the strength of the negative feedback loop influences resource decoupling efficiency, we systematically analyzed the system across a wide range of maximal flipping rates from the LR state to the BP state, with $v_{\text{off}} \in [0, 0.75]$.

**4. Mathematical Model for the System with Re-NF-FF-Controller**

In the system with a Re-NF-FF-Controller, the flipped Pbad promoter is designed to promote GFP expression. The ODE of GFP needs to be revised as follows:

$$\frac{d[GFP]}{dt} = u_{G1} * \frac{CN/Q_G}{PF_Q} + v_{G2} * \frac{(1-\phi) * CN * R_R/Q_R}{PF_Q} - d_G * [GFP]$$

The second term represents an additional expression from the flipped Pbad promoter. To simulate the tunability of the Re-NF-FF-Controller, we systematically analyzed the system across a wide range of degradation rate of excisionase ($d_X \in [0.01, 0.06]$) and the expression rate of integrase ($v_I \in [0, 0.8]$).

**5. The general Mathematical model for all the systems**

In this section, we unify all the above ODE models into a generalized framework. In the Open-Loop System, the Pbad promoter for the RFP module remains unchanged, meaning $\phi$ is fixed at 1, which can be enforced by setting $v_{\text{off}}$ = 0. To standardize the GFP expression equation across systems with different control mechanisms, we introduce a hyperparameter $\lambda$ into the second production term of GFP, allowing this term to be zero ($\lambda = 0$) for the system with only negative feedback regulation and active ($\lambda = 1$) for the system incorporating additional feedforward regulation. This allows us to formulate a single generalized model applicable to all systems,

$$\frac{d[INT]}{dt} = u_I * \frac{CN/Q_I}{PF_Q} - d_I * [INT]$$

$$\frac{d[XIS]}{dt} = u_R * \frac{CN * R_R/Q_R}{PF_Q} - d_X * [XIS]$$

$$\frac{d[AraC]}{dt} = u_A * \frac{CN/Q_A}{PF_Q} - d_A * [AraC]$$

$$\frac{d[GFP]}{dt} = u_{G1} * \frac{CN/Q_G}{PF_Q} + \lambda * u_{G2} * \frac{(1-\phi) * CN * R_R/Q_R}{PF_Q} - d_G * [GFP]$$

$$\frac{d[RFP]}{dt} = u_R * \frac{CN * \phi * R_R/Q_R}{PF_Q} - d_R * [RFP]$$

$$\frac{d[\phi]}{dt} = v_{\text{on}} * \frac{[INT]^4}{{K_{I1}}^4 + [INT]^4} * (1-\phi) - v_{\text{off}} * \frac{[XIS]^4}{{K_X}^4 + [XIS]^4} \frac{[INT]^4}{{K_{I2}}^4 + [INT]^4} * \phi$$

Where $PF_Q = 1 + CN * \left(\frac{1}{QI} + \frac{1}{Q_A} + \frac{1}{Q_G} + \frac{2*\phi*R_R}{Q_R} + \frac{(1-\phi)*R_R}{Q_R}\right), R_R = \frac{Sa*[AraC]^2}{Sa*[AraC]^2+1}, Sa = C_{min} + (C_{max} - C_{min}) * \left(\frac{Lara^n}{Lara^n + J^n}\right)$.

The parameters are, unless otherwise mentioned:

$CN = 50, v_{\text{on}} = 0.1, v_{\text{off}} = 0.5, v_A = 0.5, v_I = 0.8, v_{G1} = 0.8, v_{G2} = 2, v_R = 0.5, K_{I1} = 0.1, K_{I2} = 5, K_X = 5, J = 0.001, C_{min} = 0.000001, C_{max} = 0.015, \mathrm{n} = 3, d_A = 0.01, d_I = 0.01, d_G = 0.01, d_R = 0.01, d_X = 0.01, Q_A = 150, Q_I = 150, Q_G = 50, Q_R = 5.$

**Supplementary Table 1. BioBrick parts used in this paper.**

| Part name | Description | Biobrick number |
|---|---|---|
| J23105 | Constitutive promoter | BBa_J23105 |
| J23106 | Constitutive promoter | BBa_J23106 |
| J23110 | Constitutive promoter | BBa_J23110 |
| Ptet | Constitutive promoter | BBa_R0040 |
| Pbad | Inducible promoter activated by AraC and L-arabinose | BBa_K206000 |
| B0031 | Ribosome binding site | BBa_B0031 |
| B0034 | Ribosome binding site | BBa_B0034 |
| AraC | Arabinose operon regulatory protein | araC |
| GFP | Green fluorescent protein | BBa_E0040 |
| RFP | Red fluorescent protein | BBa_E1010 |
| CFP | Cyan fluorescent protein | BBa_E0020 |
| B0015 | Terminator | BBa_B0015 |
| B1002 | Terminator | BBa_B1002 |
| PSB1A2 | Plasmid Backbone | pSB1A2 |
| PSB3K3 | Plasmid Backbone | pSB3K3 |

**Supplementary Table 2. Sequences of parts used in this paper.**

| Part name | Description | Sequence |
|---|---|---|
| INT | Integrase | atgagagccctggtagtcatccgcctgtcccgcgtcaccgatg<br>ctacgacttcaccggagcgtcagctggagtcttgccagcagct<br>ctgcgcccagcgcggctgggacgtcgtcggggtagcggagg<br>atctggacgtctccggggcggtcgatccgttcgaccggaagc<br>gcagaccgaacctggcccggtggctagcgttcgaggagcaa<br>ccgtttgacgtgatcgtggcgtaccgggtagatcggttgacccg<br>atcgatccggcatcttcagcagctggtccactgggccgaggac<br>cacaagaagctggtcgtctccgcgaccgaagcgcacttcgat<br>acgacgacgccgtttgcggcggtcgtcatcgcgcttatgggaa<br>cggtggcgcagatggaattagaagcgatcaaagagcggaa<br>ccgttcggctgcgcatttcaatatccgcgccgggaaataccga<br>ggatccctgccgccgtggggatacctgcctacgcgcgtggac<br>ggggagtggcggctggtgccggaccctgtgcagcgagagcg<br>catcctcgaggtgtatcaccgcgtcgtcgacaaccacgagcc<br>gctgcatctggtggcccacgacctgaaccggcgtggtgtcctgt<br>cgccgaaggactacttcgcgcagctgcaaggccgcgagccg<br>cagggccgggagtggtcggctaccgcgctgaagcgatcgat<br>gatctccgaggcgatgctcgggtacgcgactctgaacggtaa<br>gaccgtccgagacgacgacggagccccgctggtgcgggctg<br>agccgatcctgacccgtgagcagctggaggcgctgcgcgcc<br>gagctcgtgaagacctcccgggcgaagcccgcggtgtctacc<br>ccgtcgctgctgctgcgggtgttgttctgcgcggtgtgcgggga<br>gcccgcgtacaagttcgccgggggaggacgtaagcacccgc<br>gctaccgctgccgctcgatggggttcccgaagcactgcggga<br>acggcacggtggcgatggccgagtgggacgcgttctgcgag<br>gagcaggtactggatctgctcggggacgcggagcgtctggag<br>aaagtctgggtagcgggctcggactccgcggtcgaactcgcg<br>gaggtgaacgcggagctggtggacctgacgtcgctgatcggc<br>tccccggcctaccgggcgggctctccgcagcgagaagcact<br>ggatgcccgtattgcggcgctggccgcgcggcaagaggagc<br>tggagggcctggaggctcgcccgtctggctgggagtggcgcg<br>agaccgggcagcggttcggggactggtggcgggagcagga<br>caccgcggcaaagaacacctggcttcggtcgatgaacgttcg<br>gctgacgttcgacgtccgcggcgggctgactcgcacgatcga<br>cttcggggatcttcaggagtacgagcagcatctcaggctcggc<br>agcgtggtcgaacggctacacaccgggatgtcgtaa(5) |
| XIS | Excisionase | atggccggcactcagcgtatcgtctttctacccgatactcagttg<br>cctttcgaggcgcgcaaagagatgcaagcggtcatccgcttc<br>atcggggatgtccagccgtacggcgtggtacatatcggtgacg<br>tcctagacctgccgcagccctcgcgctggaaccggggaacc<br>aagggcgagttcgagggctcggtgtaccgcgacgcggacta<br>cgccaagaagaacctgatggagccgctgcgcaaggtctacg<br>acggctggatcgggatgcacgagggcaaccacgatctgcga<br>gcccgcgagtacctggccaagaacgcaccggccctggagg<br>gtacgcacgctttcgacatcgacgtgctgctcgacttcgacggg<br>ttcggtgtggagctgctgcctgacttctacgacatcgctccgggc<br>tggatctccactcacgggcacatgggcaagatgacgctatccc<br>agatcgccggatcgacagcgctcaacggtgccaagaagttc |

| | | |
|---|---|---|
| | | ggcaagtccgtggtctgcggccacacgcaccggcaggctgtc gtctcgcactcgttcgggtacggcggctcggtgcgcaagaccg tcaccggcatggaagtcgggcacctgatggacatgaagaag gccaactatctaaagggcggagccgggaactggcagatggg cttcgggatgctcacggtcgacggcaagcatgtcaaggctga gatcgtcccgatcctgggaggcaagttcaccgttgacggcca ggtctgggaagtcaccggt(5) |
| attB | Attachment site attB | cggccggcttgtcgacgacggcgcactccgtcgtcaggatcat ccgggc(6) |
| attP | Attachment site attP | gtcgtggtttgtctggtcaaccaccgcgcactcagtggtgtacg gtacaaaccccgac(6) |
| attL | Attachment site attL | cggccggcttgtcgacgacggcgcactcagtggtgtacggta caaaccccgac |
| attR | Attachment site attR | gtcgtggtttgtctggtcaaccaccgcgcactccgtcgtcaggat catccgggc |
| attR-M | Mutated attachment site attR | gtcgtggtttgtctggtcaaccaccgcggtctccgtcgtcaggat catccgggc |
| AAK-tag | Fast degradation tag | gcagcaaacgacgaaaactacgctgctgctaaa |
| LVA-tag | Fast degradation tag | gctgcaaacgacgaaaactacgctttagtagct |
| Spacer | Spacer | agatggttagtgtgcgtatctgatacttttatgtctgtcctacaactc caagcgtacagttcaggccaacaccctagctttgagaaagtc gcgtcagtaaatgtcgtttccctaaagcttacaatttacctcctcg ctatt |

**Supplementary Table 3. List of circuit assembly.**

| Circuit name | Circuit design* |
| --- | --- |
| C5 | J23110 + B0034 + AraC + B0015 + B0015 (R) + RFP-Lva (R) + B0034 (R) + attB + B0015 (R) + Pbad + attp (R) + B0034 + GFP-Lva + B0015 + PSB1A2 |
| C6 | J23110 + B0034 + AraC + B0015 + B0015 (R) + RFP-Lva (R) + B0034 (R) + attB + B0015 (R) + Pbad + attp (R) + B0034 + INT + B0034 + GFP-Lva + B0015 + PSB1A2 |
| C21 | J23110 + B0034 + AraC + B0015 + B1002 (R) + XIS (R) + B0034 (R) + RFP (R) + B0034 (R) + attL + Pbad (R) + B0015 + attR (R) + B0034 + GFP + B0015 + PSB1A2 |
| C24 | J23110 + B0034 + AraC + B0015 + B1002 (R) + XIS (R) + B0034 (R) + RFP (R) + B0034 (R) + attL + Pbad (R) + B0015 + attR (R) + B0034 + GFP + B0015 + J23105 + B0031 + INT + B0015 + PSB1A2 |
| C25 | J23110 + B0034 + AraC + B0015 + B1002 (R) + XIS (R) + B0034 (R) + RFP (R) + B0034 (R) + attL + Pbad (R) + B0015 + attR (R) + B0034 + GFP + B0015 + J23105 + B0031 + INT (start codon ATG to GTG) + B0015 + PSB1A2 |
| C34 | B1002 (R) + RFP (R) + B0034 (R) + attL + Pbad (R) + B0015 + attR (R) + B0015 + J23106 + B0034 + GFP + B0015 + J23110 + B0034 + AraC + B0015 + PSB3K3 |
| C35 | B1002 (R) + XIS (R) + B0034 (R) + RFP (R) + B0034 (R) + attL + Pbad (R) + B0015 + attR (R) + B0015 + J23106 + B0034 + GFP + B0015 + J23110 + B0034 + AraC + B0015 + J23105 + B0031 + INT + B0015 + PSB3K3 |
| C37 | B1002 (R) + AAK-tag (R) + XIS (R) + B0034 (R) + RFP (R) + B0034 (R) + attL + Pbad (R) + B0015 + attR (R) + B0015 + J23106 + B0034 + GFP + B0015 + J23110 + B0034 + AraC + B0015 + J23105 + B0031 + INT + B0015 + PSB3K3 |
| C42 | B1002 (R) + RFP (R) + B0034 (R) + attL + Spacer (R) + Pbad (R) + B0015 + attR (R) + B0015 + J23106 + B0034 + GFP + B0015 + J23110 + B0034 + AraC + B0015 + PSB3K3 |
| C43 | B1002 (R) + XIS (R) + B0034 (R) + RFP (R) + B0034 (R) + attL + Spacer (R) + Pbad (R) + B0015 + attR (R) + B0015 + J23106 + B0034 + GFP + B0015 + J23110 + B0034 + AraC + B0015 + J23105 + B0031 + INT + B0015 + PSB3K3 |
| C44 | J23106 + B0034 + GFP + B0015 + J23110 + B0034 + AraC + B0015 + J23105 + B0031 + INT + B0015 + B1002 (R) + XIS (R) + B0034 (R) + RFP (R) + B0034 (R) + attL + Spacer (R) + Pbad (R) + B0015 + attR (R) + B0015 + PSB3K3 |
| C45 | B1002 (R) + RFP (R) + B0034 (R) + attL + Spacer (R) + Pbad (R) + B0015 + attR (R) + B0015 + J23106 + B0034 + GFP + B0015 + J23110 + B0034 + AraC + B0015 + Ptet + B0034 + CFP + B0015 + PSB3K3 |
| C46 | B1002 (R) + XIS (R) + B0034 (R) + RFP (R) + B0034 (R) + attL + Spacer (R) + Pbad (R) + B0015 + attR (R) + B0015 + J23106 + B0034 + GFP + B0015 + J23110 + B0034 + AraC + B0015 + J23105 + B0031 + INT + B0015 + Ptet + B0034 + CFP + B0015 + PSB3K3 |
| C52 | J23106 + B0034 + GFP + B0015 + J23110 + B0034 + AraC + B0015 + J23105 + B0031 + INT + B0015 + B1002 (R) + XIS (R) + B0034 (R) + RFP (R) + B0034 (R) + attL + Spacer (R) + Pbad (R) + B0015 + attR-M (R) + B0015 + PSB3K3 |

| | |
|---|---|
| C53 | B1002 (R) + XIS (R) + B0034 (R) + RFP (R) + B0034 (R) + attL + Spacer (R) + Pbad (R) + B0015 + attR (R) + J23106 + B0034 + GFP + B0015 + J23110 + B0034 + AraC + B0015 + J23105 + B0031 + INT + B0015 + PSB3K3 |
| C55 | B1002 (R) + XIS (R) + B0034 (R) + RFP (R) + B0034 (R) + attL + Spacer (R) + Pbad (R) + B0015 + attR-M (R) + J23106 + B0034 + GFP + B0015 + J23110 + B0034 + AraC + B0015 + J23105 + B0031 + INT + B0015 + PSB3K3 |
| C57 | B1002 (R) + XIS (R) + B0034 (R) + RFP (R) + B0034 (R) + attL + Spacer (R) + Pbad (R) + B0015 + attR (R) + J23106 + B0034 + GFP + B0015 + J23110 + B0034 + AraC + B0015 + J23105 + B0031 + INT (start codon ATG to GTG) + B0015 + PSB3K3 |
| C59 | B1002 (R) + AAK-tag (R) + XIS (R) + B0034 (R) + RFP (R) + B0034 (R) + attL + Spacer (R) + Pbad (R) + B0015 + attR (R) + J23106 + B0034 + GFP + B0015 + J23110 + B0034 + AraC + B0015 + J23105 + B0031 + INT + B0015 + PSB3K3 |
| C65 | attB + B0015 (R) + Pbad + Spacer + attP (R) + B0034 + GFP + B0015 + J23110 + B0034 + AraC + B0015 + PSB3K3 |
| C66 | attB + B0015 (R) + Pbad + Spacer + attP (R) + B0015 + B0034 + GFP + B0015 + J23110 + B0034 + AraC + B0015 + PSB3K3 |
| C70 | attB + B0015 (R) + Pbad + Spacer + attP (R) + B0015 + B1002 + B0034 + GFP + B0015 + J23110 + B0034 + AraC + B0015 + PSB3K3 |
| C73 | B1002 (R) + RFP (R) + B0034 (R) + attL + Spacer (R) + Pbad (R) + B0015 + attR (R) + B0015 + B1002 + J23106 + B0034 + GFP + B0015 + J23110 + B0034 + AraC + B0015 + PSB3K3 |
| C74 | B1002 (R) + XIS (R) + B0034 (R) + RFP (R) + B0034 (R) + attL + Spacer (R) + Pbad (R) + B0015 + attR (R) + B0015 + B1002 + J23106 + B0034 + GFP + B0015 + J23110 + B0034 + AraC + B0015 + J23105 + B0031 + INT + B0015 + PSB3K3 |

* (R) indicates that the gene is arranged in reverse orientation within the circuit.
